# Intrinsic bursts facilitate learning of Lévy flight movements in recurrent neural network models

**DOI:** 10.1101/2021.11.15.468754

**Authors:** Morihiro Ohta, Toshitake Asabuki, Tomoki Fukai

## Abstract

Isolated spikes and bursts of spikes are thought to provide the two major modes of information coding by neurons. Bursts are known to be crucial for fundamental processes between neuron pairs, such as neuronal communications and synaptic plasticity. Deficits in neuronal bursting can also impair higher cognitive functions and cause mental disorders. Despite these findings on the roles of bursts, whether and how bursts have an advantage over isolated spikes in the network-level computation remains elusive. Here, we demonstrate in a computational model that not isolated spikes but intrinsic bursts can greatly facilitate learning of Lévy flight random walk trajectories by synchronizing burst onsets across neural population. Lévy flight is a hallmark of optimal search strategies and appears in cognitive behaviors such as saccadic eye movements and memory retrieval. Our results suggest that bursting is a crucial component of sequence learning by recurrent neural networks in the brain.

## INTRODUCTION

Neurons in the brain display variety of temporal discharging patterns, among which bursting represents the generation of multiple spikes with brief inter-spike intervals (typically, several milliseconds) in a short period of time (typically, several tens to hundreds of milliseconds). Bursting neurons are found ubiquitously in the brain and are thought to play active roles in transferring and routing information [1–5], inducing synaptic plasticity [6, 7], and supporting and/or altering cognitive functions [2, 7–14]. Deficits in burst generation can cause mental disorders [15, 16]. While our understanding of the roles of bursting has been advanced, the computational advantages of spike bursts over isolated spikes remain elusive.

Here, we show the benefits of bursting activity in learning sequences generated by a special class of random walks observed in various animal behaviors. We investigate whether and how bursting neurons improve the ability of neural network models to learn the dynamical trajectories of Lévy flight, which is a random walk with step sizes obeying a heavy-tailed distribution [17–19]. As a consequence, Lévy flight consists of many short steps and rare long-distance jumps. A well-known characteristic of Lévy flight is that it makes search more efficient than Brownian walks which only consist of relatively short steps [20, 21]. Many processes observed in biology [22–24] and physics [25, 26] can be described as Lévy flight. In neuroscience, an interesting example of Lévy flight is the stochastic trajectories of saccadic eye movement [27] on which the visual exploration of the objects of interest significantly relies. Several cortical and subcortical regions including the frontal eye field, superior colliculus, and cerebellar cortex participate in controlling and executing saccades [28] and various neurons show spike bursts in these regions [8, 9, 29, 30]. Other examples of Lévy flight are found in memory processing of animals. In the spatial exploration of rodents, the animal spends the majority of time for exploring small local areas but occasionally travels to distant places at greater speeds [31]. Hippocampal [10] and subicular [32] neurons can learn spatial receptive fields and are known to exhibit burst firing. In human subjects, memory recall can be viewed as foraging behavior obeying Lévy flight [33–35]. The appearance of Lévy flight in various types of foraging behavior and the participation of bursting neurons in the relevant brain regions motivate us to explore what benefits neuronal bursting brings to the learning and execution of such behavior.

For this purpose, we employ reservoir computing (RC) that uses a recurrent network model and FORCE learning of information-readout neurons for efficient learning of time-varying external signals (i.e., teaching signals) [36]. Originally, RC and FORCE learning were formulated for rate-coding neurons, and FORCE learning of continuous dynamical trajectories is generally fast. The RC system was also quite successful in modeling neural activities recorded from various cortical areas [37–40]. Later, RC was extended to networks of spiking neurons [41, 42], and variants of FORCE learning or some other learning method [43] for spiking neurons have also been proposed [44–46]. Results of the previous studies have indicated that isolated spikes are sufficient for learning smooth trajectories. However, whether and how isolated spikes and bursts contribute differently to learning a more general class of sequences has not been explored. In this study, we clarify this by using a spiking-neuron version of FORCE learning for training an RC system of bursting neurons.

## RESULTS

### Network model

Our model follows the conventional framework of reservoir computing except that neurons constituting a recurrent network called reservoir have regular-spiking (RS) and bursting modes (Fig. 1a). In the RS mode, the neurons tend to generate isolates spikes (Fig. 1b) whereas they are strongly bursty in the bursting mode (Fig. 1c). Neurons in the reservoir project to two readout neurons to describe the two-dimensional coordinates (*x*_1_*, x*_2_) of Lévy flight, and the outputs of these neurons are fed back to all neurons in the reservoir. We describe neurons in the reservoir with the Izhikevich model, which is able to mimic the temporal discharging patterns of various neurons [47]:

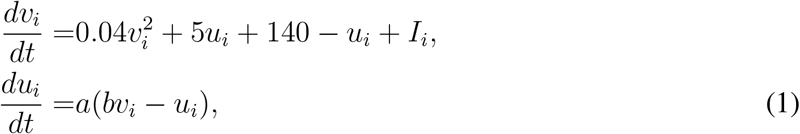

where *a* = 0.02 and *b* = 0.2, and *i* is a neuron index. We set as *c* = −65 mV and *d* = 8 in the RS mode and *c* = −50 mV and *d* = 2 in the bursting mode. The values of *v_i_* and *u_i_* are reset to *c* and *u_i_* + *d* when *v_i_* reaches the threshold of 30 mV. We use this model for simplicity of numerical simulations although the Izhikevich model does not take refractory periods into account and may exhibit unrealistically high frequency bursting.

**Fig. 1.**
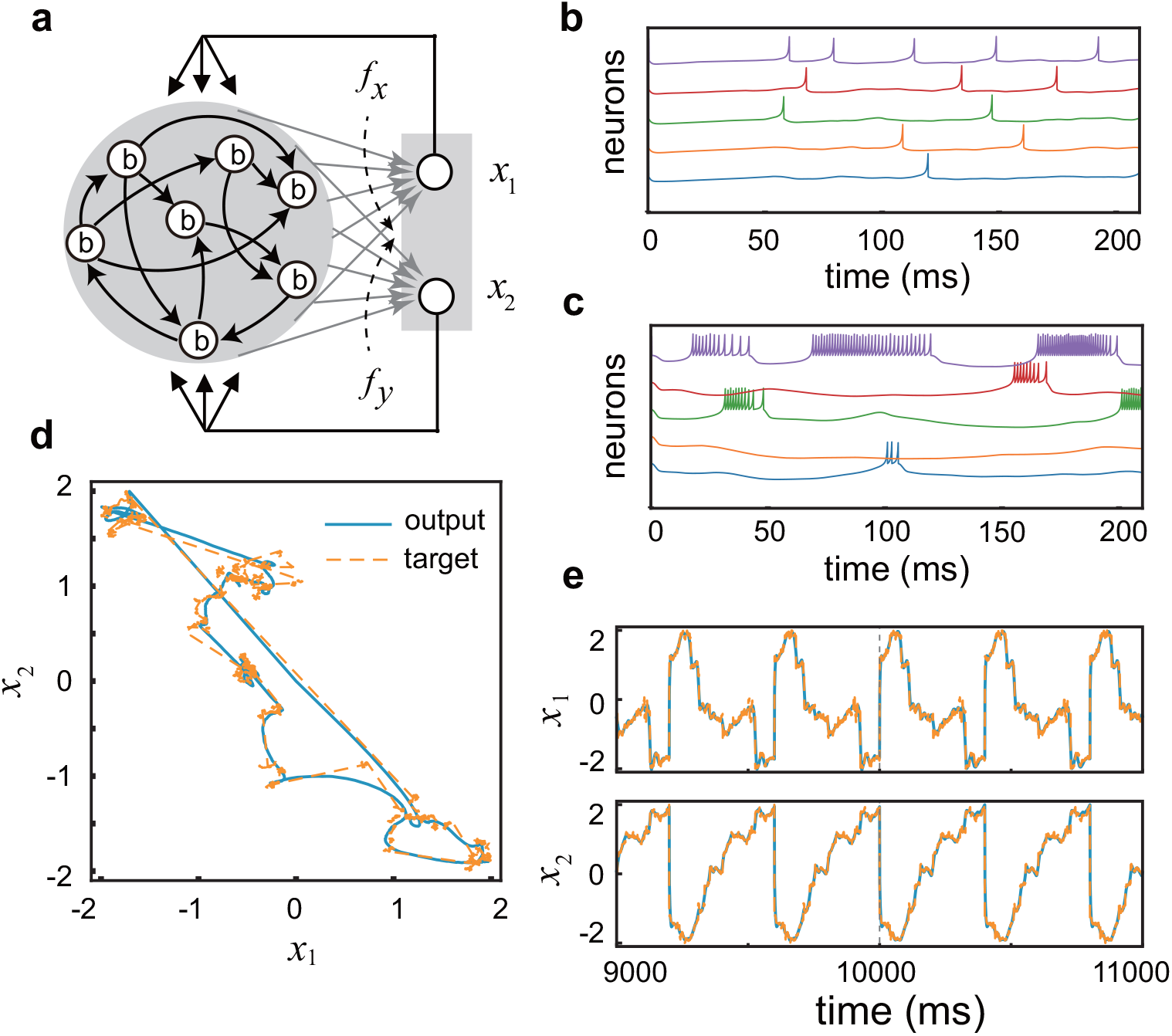
The architecture and basic performance of the model. (a) The present RC system consists of a reservoir and two readout neurons. (b, c) Before learning, neurons in the reservoir tend to show isolated spikes in the RS mode (b) or intrinsic bursts in the bursting mode (c). Here, the firing patterns were simulated at *G* = 50. (d) A typical example of the target trajectories representing a finite portion of two-dimensional Lévy flight (orange) and the learned responses of the readout neurons (blue). (e) The time evolution of the two readout neurons are shown as functions of time. Large discontinuous jumps in (d) and (e) indicate the onset and end point of the repeated target signal.

Synaptic current is given as *I_i_* = *s_i_*(*t*) + *I_b_*, where *I_b_* is a constant bias and recurrent synaptic inputs are

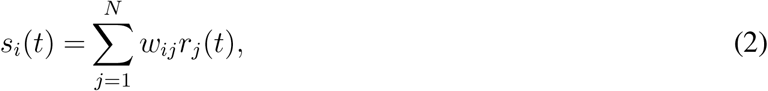

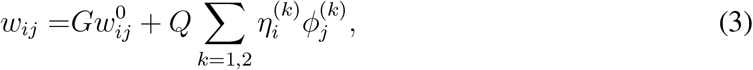

in terms of the instantaneous firing rate *r_i_*(*t*) of neuron *i* at time *t*. The synaptic weight matrix *w_ij_* has non-modifiable components 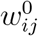 and modifiable components 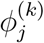, with *G* and *Q* being constant parameters. While *Q* = 100 throughout this paper, the value of *G* is mode-dependent, as shown later. The encoding parameter 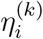 is randomly drawn from the uniform distribution [−1, +1]^*k*^, where *k* = 2 for the target trajectories representing a two-dimensional Lévy flight. The linear decoder 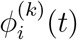 determines activities of the readout units *x*^(*k*)^(*t*):

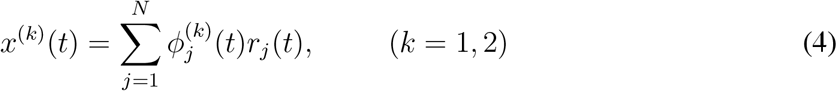

which should approximate a given target trajectory. See Methods for the details of construction of Lévy flight and FORCE learning.

### Advantages of bursts over isolated spikes in sequence learning

During learning, the model was repeatedly exposed to a periodic target signal representing the repetition of a finite portion of Lévy flight trajectories (Methods). The model can learn these trajectories in either RS or bursting mode. Large jumps in the trajectory are thought to be difficult for the model to accurately learn. As we will show later, the accuracy and speed of learning significantly depend on the mode of firing. Figure 1d displays an example of the time-varying output of the two readout neurons after the model learned a target signal in the bursting mode. As expected, the output of the model tends to deviate largely from the target trajectory when it shows relatively large jumps. Nonetheless, overall the model well replicates the target trajectory in the burst mode even after the learning process is turned off. The agreement between the target trajectory and the model’s output is more clearly visible in the time evolution of the variables *x*_1_ and *x*_2_ (Fig. 1e).

We quantitatively compare the performance of the model in learning between the bursting and RS modes. The strength of synaptic connections that gives an optimal performance may differ in the individual modes. To make a fair comparison, we first search an optimal coupling strength that minimizes the error in each mode. We calculate the average errors between a target trajectory and an actual output in the bursting mode and the RS mode as a function of the connection strength *G*. Figure 2a and 2b show the errors obtained after 25 and 50 trials of learning, respectively, when the target length is 400 ms. For each value of *G*, the standard deviations of the error are calculated over simulations with 20 different initial conditions. As we can see from these figures, the error is minimized for relatively weak connections (*G* ~ 50) in the bursting mode. In contrast, the model achieves the least error at much stronger connections (*G* ~ 170) in the RS mode. The minimum average error is slightly smaller in the bursting mode than in the RS mode although the error sizes are not greatly different between the two modes after 50 cycles of training (Fig. 2b). Given these results, one might conclude that spike bursts have little advantage over isolated spikes in the present sequence learning task.

**Fig. 2.**
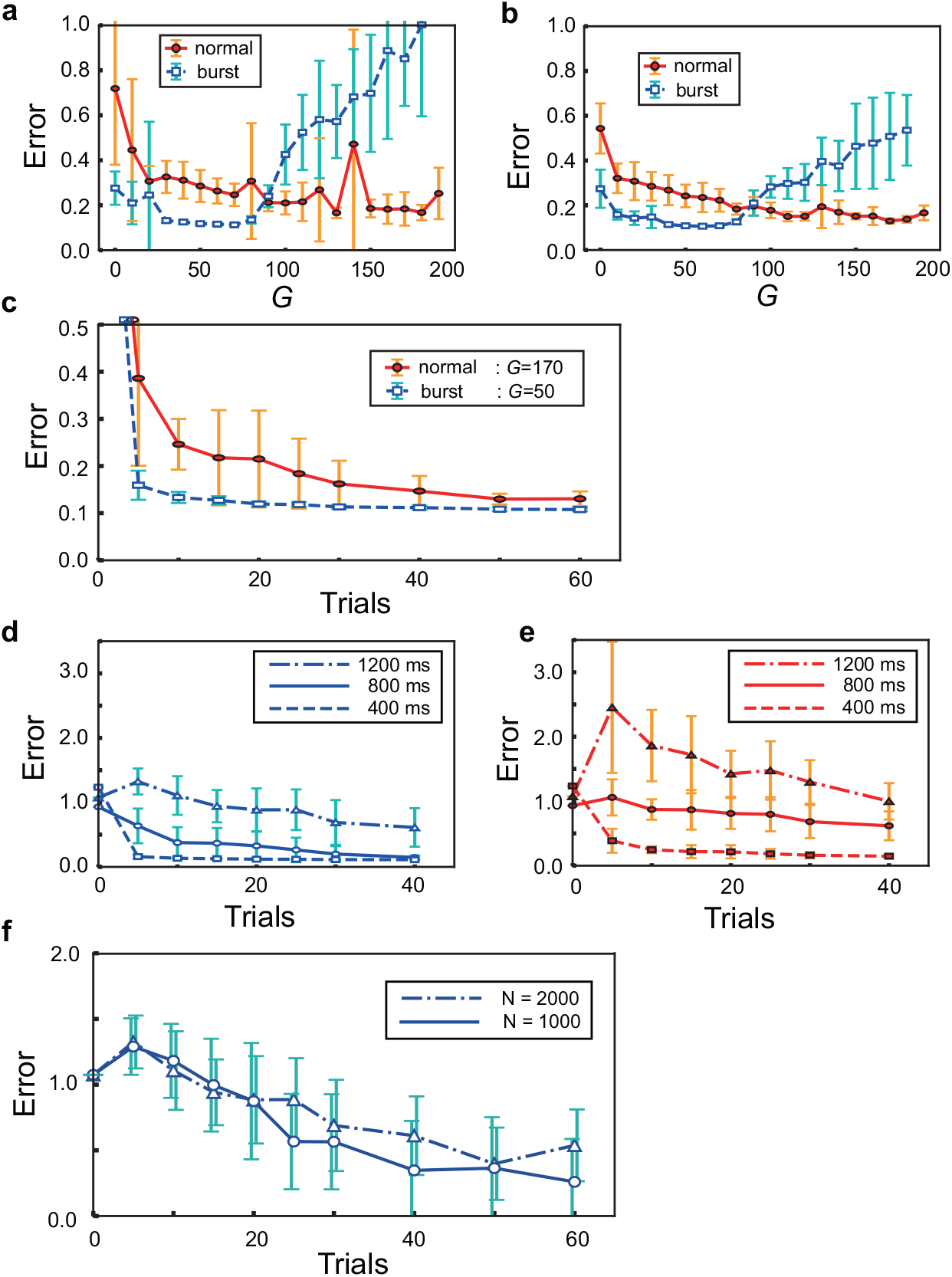
Learning in the burst vs. RS modes. (a) Errors in the bursting and RS modes are plotted against the strength of recurrent connections after 25 learning trials. Error bars show the standard deviations. (b) Similar errors are plotted after 50 learning trials. (c) The time courses of errors during learning are shown for the optimal coupling strengths of the individual modes. (d, e) Similar time courses are shown in the bursting (d) and RS (e) modes for target signals of lengths 400, 800, and 1200 ms. The plots for 400 ms are copied from (c). (f) Error time courses are shown in the bursting mode when the target length is 1200 ms and the reservoir size is 1000 or 2000.

However, the results presented in Fig. 2a and 2b reveal two intriguing differences in learning between the RS mode and the bursting mode. First, while the two modes yield approximately the same minimum values of average errors, the bursting mode yields a much smaller variance at the minimum error than the RS mode. In particular, Figure 2a demonstrates that the variance almost vanishes for the optimal range of *G* values after 25 training cycles in the bursting mode. This is not the case for the optimal range of *G* values in the RS mode. Second and more importantly, the average error decreases much faster during learning in the bursting mode than in the RS mode, showing impressively different learning speeds between the two modes (Fig. 2c). Generally, the FORCE learning enables rapid learning of a smooth target trajectory even if the trajectory is chaotic [36]. However, our results show that the FORCE learning with isolated spikes requires several tens of trials for learning a target trajectory representing random walks of Lévy flight. In strong contrast, spike bursts enable the same rule to learn such a target trajectory at a similar accuracy within only ten trials. The merits of bursting are also suggested by the common observation that the individual neurons tend to generate spike bursts after learning at the corresponding optimal coupling strength irrespective of the mode (Supplementary Fig. 1).

As the length of target trajectories is increased, performance in sequence learning is degraded in both modes. However, the superiority of the bursting mode over the RS mode in rapid sequence learning remains hold (Fig. 2d, e). We note that the absolute values of errors are not really meaningful. These values become smaller as we include more neurons in the reservoir (Fig. 2f).

### Learning through burst synchronization

Now, we investigate why and how spike bursts improve the performance of the network model in learning trajectories of Lévy flight. We show that synchronized bursting of neurons plays an active role in the present sequence learning. Figure 3a shows the time evolution of a portion of the learned trajectory *x*_1_(*t*) and *x*_2_(*t*) with vertical dashed lines indicating the times of long-distance flights. Here, a long-distance flight refers to a step (Δ*x*_1_, Δ*x*_2_) of which the length 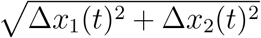 is greater than 0.16, which approximately corresponds to the top 5% of long-distance jumps. In Fig. 3b and 3c, we show spike raster of 100 bursting neurons chosen randomly from the reservoir during the corresponding period of time before and after learning, respectively. While there are many neurons that rarely fire, some neurons intermittently generate brief (~ 30 ms) to prolonged (~ 150 ms) high-frequency bursts. The individual neurons change their firing patterns before and after learning, but the distributions of inter-spike intervals at the population level remain almost unchanged during learning (Supplementary Fig. 2a, b).

**Fig. 3.**
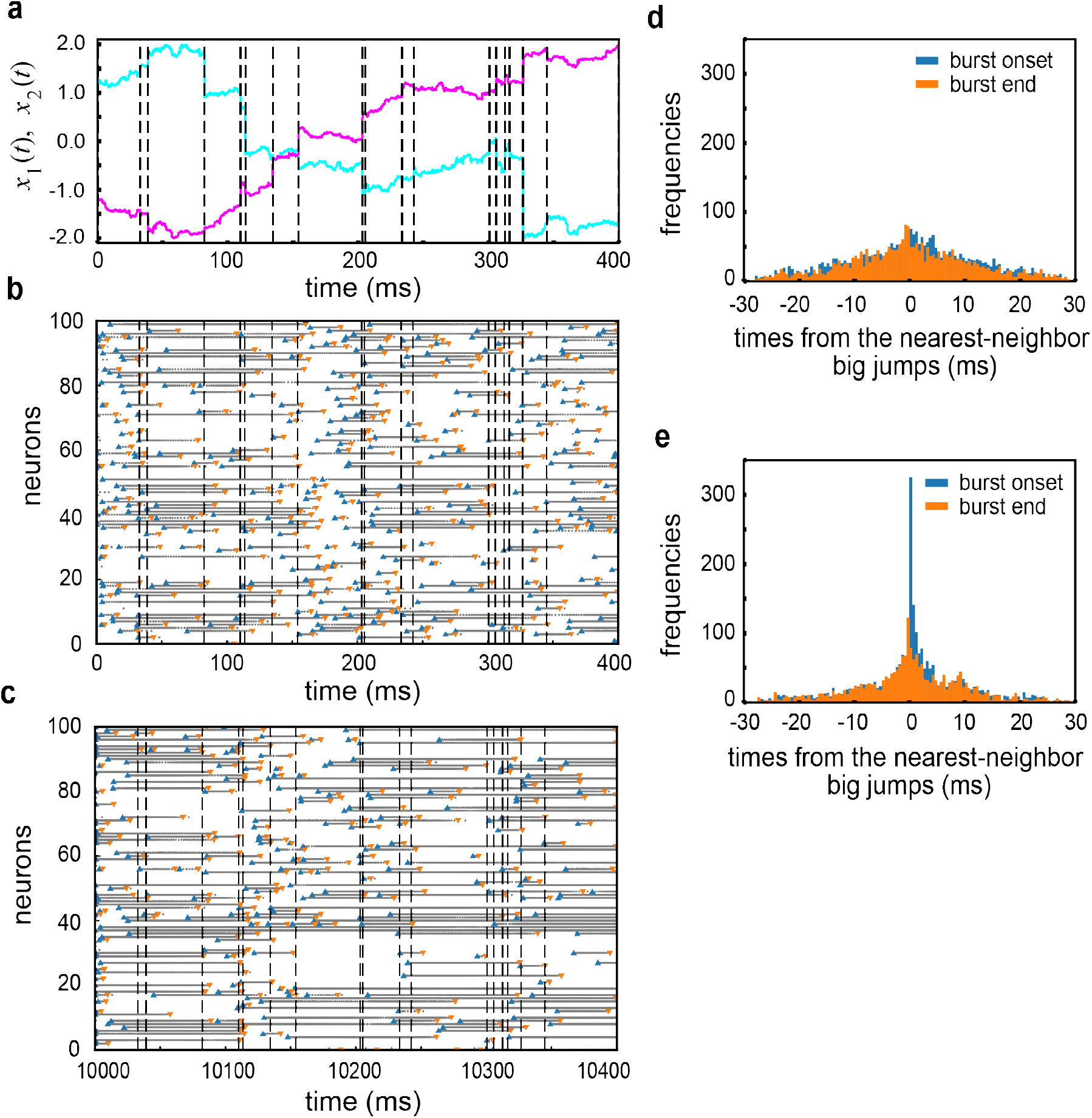
Temporal coordination of bursts by learning. (a) A two-dimensional target trajectory shows big jumps at the times indicated by vertical dashed lines. (b) Spike raster of 100 neurons sampled randomly from the reservoir before learning. (c) Spike raster is shown for the same neurons after learning. (d, e) Distributions of the onset and end times of bursts around the times of big jumps are calculated before (d) and after (e) learning.

However, visual inspection of the spike raster suggests that many neurons start or stop generating spike bursts around the times of large flights after learning and that such a tendency is weak before learning. Therefore, regarding that spikes with their inter-spike intervals shorter than 3 ms belonged to a burst, we identified the onsets and end times of bursts of individual neurons and calculated the distributions of the onset/end times of bursts relative to the times of the nearest large jumps (i.e., the times of burst onsets/ends minus the times of the nearest neighbor large flights) before (Fig. 3d) and after learning (Fig. 3e). Intriguingly, the post-learning distributions exhibited sharp peaks around the origin of the axis for the relative time. The relative times of burst onsets show a particularly prominent peak. These results reveal that the RC system operating in the bursting mode learns the target trajectory of Lévy flight by shifting the times of bursts close to the occurrence times of large jumps. In other words, the RC system synchronizes bursting of the individual neurons around the times of large jumps. This synchronization of bursts is thought to advantage recurrent networks of bursting neurons in learning of sequences that involve abrupt changes in the trajectories.

## DISCUSSION

We have trained an RC system of spiking neurons on a difficult sequence learning task where the target sequence represents random walks. FORCE learning can project the neural population activity of the reservoir quickly onto a target trajectory for a wide range of continuous trajectories including chaotic ones. This fast convergence of learning is a merit of RC, making RC useful for various practical applications. However, when a target trajectory consists of abrupt steps including long-distance jumps, as was the case in Lévy flight, FORCE learning with isolated spikes requires a large number of trials for minimizing the error signal. In contrast, the same learning rule can rapidly minimize the error by aligning the onsets as well as the end times of bursts in the neighborhoods of the times of long-distance jumps. This implies that the system synchronizes bursts of the individual neurons around these times. Thus, the RC system can learn the Lévy flight trajectories much faster with bursts than with isolated spikes. Our model suggests that bursts contribute crucially to learning foraging-like cognitive behaviors.

Our results show an interesting qualitative agreement with some experimental observations. It has been known that the onsets of bursts in the saccade-related burst neurons are tightly linked to saccade onsets in the superior colliculus [8, 9]. These neurons tend to discharge prior to a saccade if the movement is in their preferred direction, and their discharges follow rather than precede saccades for movements deviating from their preferred directions. Altough our model is far simpler compared to the actual neural circuits that control saccadic eye movements [48], the sharp peak of burst onsets around the times of long-distance steps in Fig. 3e seems to be consistent with the characteristic behavioral correlates of the saccade-related burst neurons in the superior colliculus.

During spatial navigation, hippocampal place cells exhibit both bursts and isolated spikes [3], and the different discharging patterns are thought to play distinct functional roles in the hippocampal memory processing [3, 11, 49]. The hippocampal area CA3, which has prominent recurrent excitatory connections, resembles a reservoir in this model. Furthermore, an abstract model of the entorhinal-hippocampal memory system accounted for the different statistical structures of hippocampal sequence generation, such as diffusive vs. Lévy flight-like random walks [31]. Therefore, the hippocampal circuits are of potential relevance to this study. However, the relationships between spatial information coding and the cells’ discharging patterns are not simple, depending on specific cell types and brain regions [32, 49]. To our knowledge, whether CA3 neural population synchronizes their burst discharges around the times of long-distance runs of animals has not been known. On the other hand, it is known that bursts of CA3 neurons mostly occur in an inbound travel towards their receptive field centers [10]. Clarifying the distinct computational roles of isolated spikes and bursts to the hippocampal memory processing is an intriguing open question.

In summary, this study showed the advantages of bursting neuronal activity in rapid learning of dynamical trajectories obeying Lévy flight. Bursting is ubiquitously found in various regions of the brain, and previous studies suggest the active roles of bursts in robust spike propagation and induction of synaptic plasticity. Our results give a further insight into the unique role of bursts at the network-level learning and computation.

## METHODS

### Lévy flight

Trajectories obeying Lévy flight were generated by using the function, scipy.stats.levy_stable.rvs(), in the Scipy library of Python for scientific calculations. This function generates a series of random numbers that obey the Lévy distribution [17, 18]. In short, a Lévy stable distribution has the characteristic function of the form,

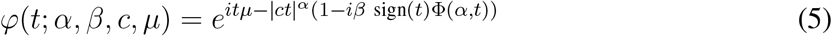

where *α*, *β*, *c*, and *μ* are the characteristic exponent, skewness parameter, scale parameter, and location parameter, respectively, and

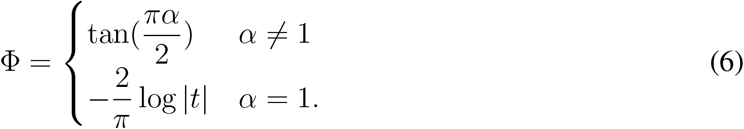

The probability density function for Lévy stable distributions is given as

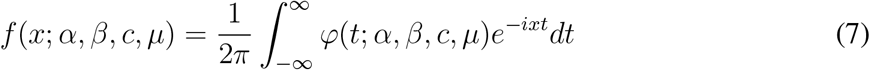

where −∞ < *x* < ∞. Throughout this study, we set as *α* = 1.5, *β* = 0, *c* = 1 and *μ* = 0.

Now, step sizes of a two-dimensional Lévy walk can be written as

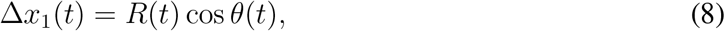

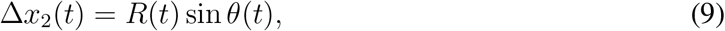

where the angle of each step *θ*(*t*) is drawn randomly from the uniform distribution 0 ≤ *θ* ≤ 2*π*, and the step amplitude *R*(*t*) was determined as *R* = *F*^−1^(*r*), where

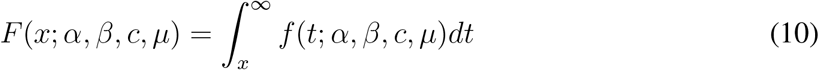

is the cumulative distribution function of *f* (*x*; *α, β, c, μ*) and 0 < *r* ≤ 1 is a uniform random number.

We limited the target trajectories with in a square area |*x*_1_| ≤ 2, |*x*_2_| ≤ 2 by normalizing the coordinates of Lévy walk as

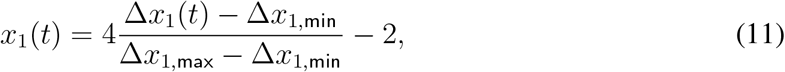

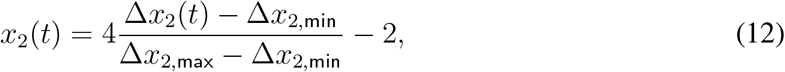

where Δ*x*_*k*,min_ and Δ*x*_*k*,max_ (*k* = 1, 2) stand for the minimum and maximum values of the previous and current step sizes Δ*x*_*k*_(*t*′) (*t*′ ≤ *t*), respectively.

### FORCE learning

We used a straight-forward extension of the FORCE learning to spiking neurons [46]. A double exponential filter was used to low-pass filter the individual spikes of the *i*-th neuron in the reservoir:

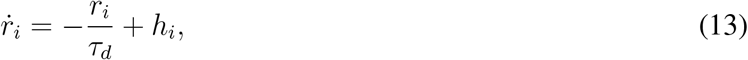

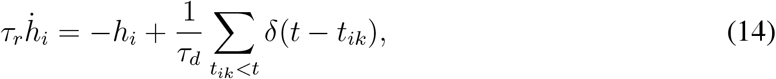

where *τ_r_* and *τ_d_* are the synaptic rise time and synaptic decay time, respectively. Values of these parameters were set as *τ_r_* = 2 ms and *τ_d_* = 20 ms.

Using the error signals *e*^(*k*)^(*t*) = *f*^(*k*)^(*t*) − *x*^(*k*)^(*t*), we update the decoders as follows:

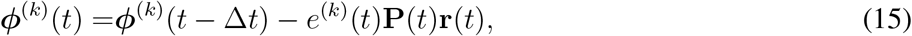

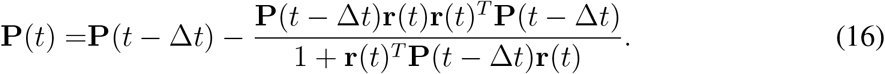

The initial conditions are given as 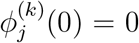 and **P**(0) = **I**_*N*_ /λ, where **I**_*N*_ is an *N* -dimensional identity matrix and *λ* = 10 for both regular and bursting modes.

## ACKNOWLEDGMENTS

This work was partly supported by Grants-in-Aid for Specially Promoted Research (JSPS KAKENHI) no. 18H05213.

---

**Supplementary Figure 1.**
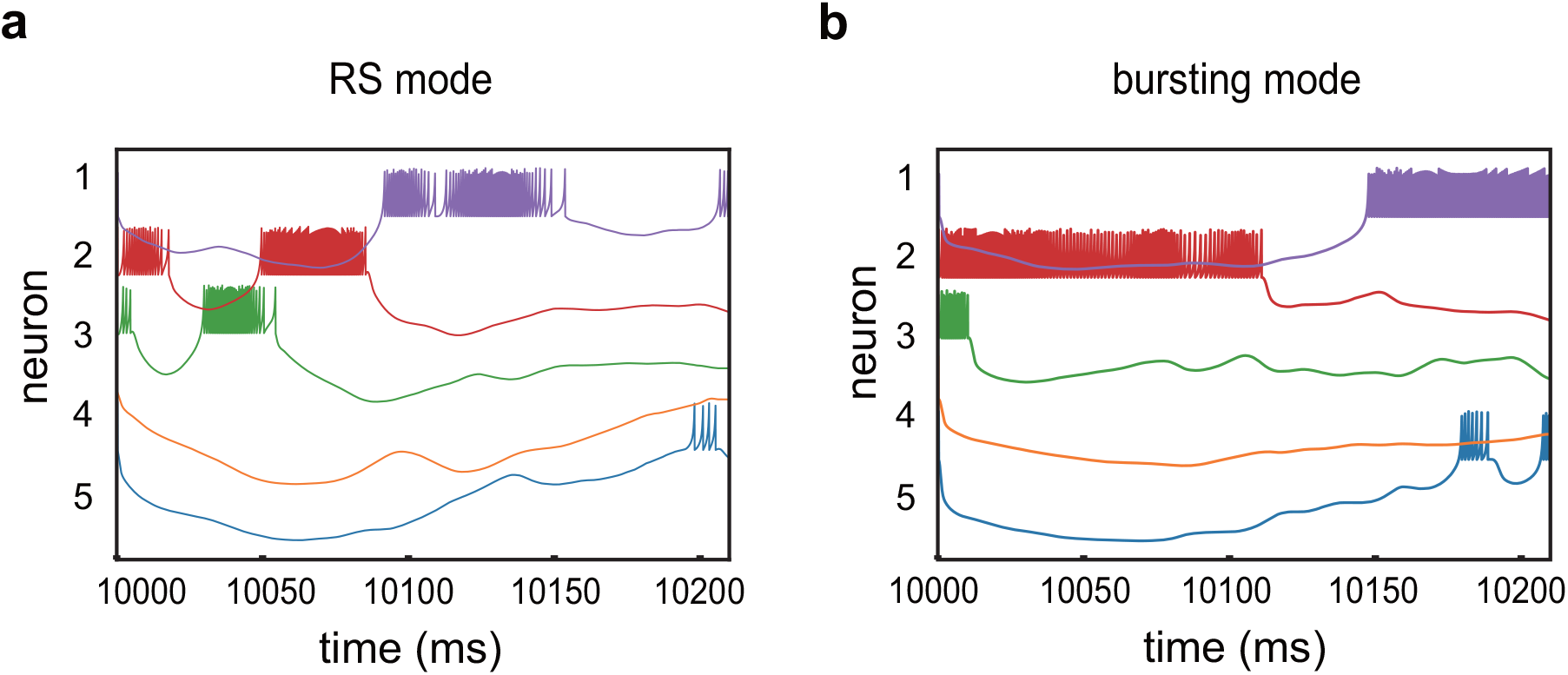
Post-learning firing patterns. (a, b) Temporal spiking patterns after learning in the RS mode (a) or bursting mode (b) are plotted for five neurons. These patterns were obtained at the optimal coupling strengths of the individual modes.

**Supplementary Figure 2.**
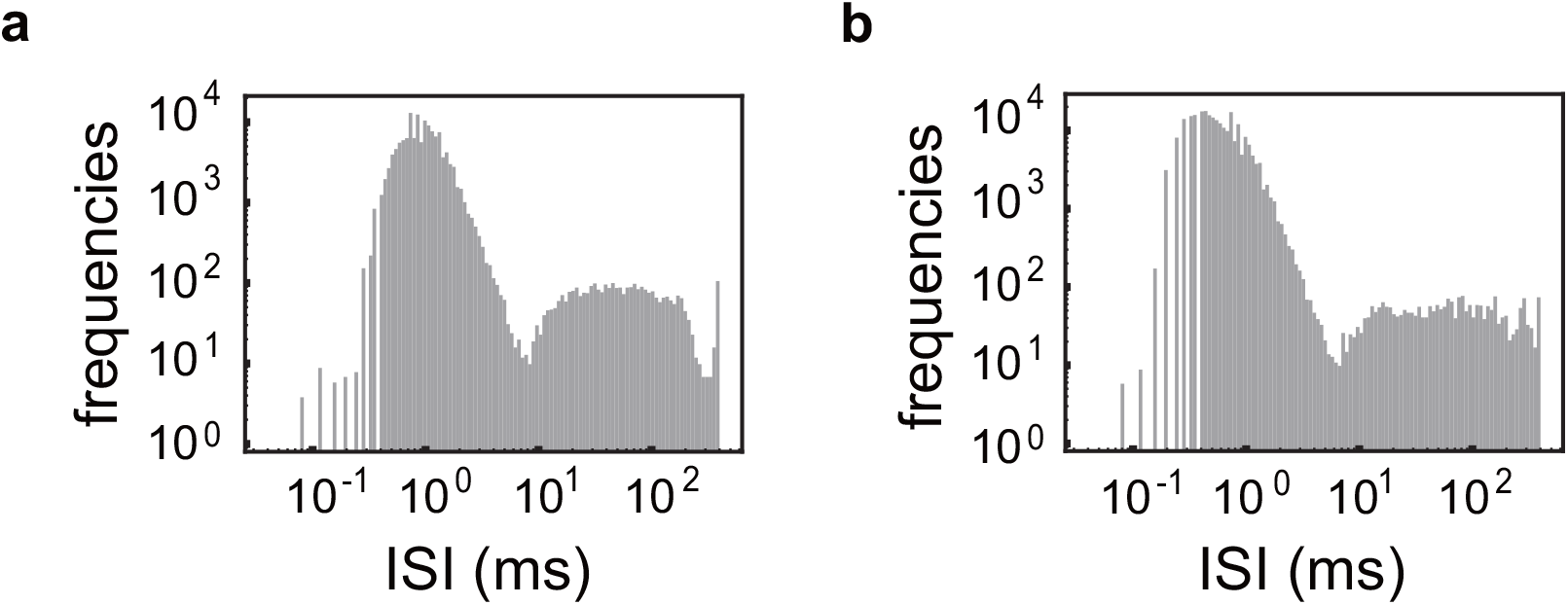
The post-learning inter-spike-interval distributions. (a, b) The inter-spike-interval distributions are calculated over all neurons in the reservoir after learning in the RS mode (a) and bursting mode (b). The Izhikevich model used in this study does not take refractory periods into account and occasionally generates unrealistically short ISIs. Sharp upper bounds at 400 ms represent the length of the target signals used in the simulations.

## REFERENCES

[1] Lisman, J. E. Bursts as a unit of neural information: making unreliable synapses reliable. Trends. Neurosci. 20, 38–43 (1997).

[2] Fanselow, E. E., Sameshima, K., Baccala, L. A. & Nicolelis, M. A. Thalamic bursting in rats during different awake behavioral states. Proc. Natl. Acad. Sci. USA 98, 15330–15335 (2001).

[3] Harris, K. D., Hirase, H., Leinekugel, X., Henze, D. A. & Buzsáki, G. Temporal interaction between single spikes and complex spike bursts in hippocampal pyramidal cells. Neuron 32, 141–149 (2001).

[4] Izhikevich, E. M., Desai, N. S., Walcott, E. C. & Hoppensteadt, F. C. Bursts as a unit of neural information: selective communication via resonance. Trends. Neurosci. 26, 161–167 (2003).

[5] Naud, R. & Sprekeler, H. Sparse bursts optimize information transmission in a multiplexed neural code. Proc. Natl. Acad. Sci. USA 115, E6329–E6338 (2018).

[6] Larson, J. & Lynch, G. Induction of synaptic potentiation in hippocampus by patterned stimulation involves two events. Science 232, 985–988 (1986).

[7] Yin, L. et al. Autapses enhance bursting and coincidence detection in neocortical pyramidal cells. Nat. Commun. 9, 4890 (2018).

[8] Goossens, H. H. L. M. & van Opstal A. J. Optimal Control of Saccades by Spatial-Temporal Activity Patterns in the Monkey Superior Colliculus. PLoS Comput. Biol. 8, e1002508 (2012).

[9] Sparks, D. L. & Mays, L. E. Movement fields of saccade-related burst neurons in the monkey superior colliculus. Brain Res. 190, 39–50 (1980).

[10] Mizuseki, K., Royer, S., Diba, K. & Buzsaki, G. Activity dynamics and behavioral correlates of CA3 and CA1 hippocampal pyramidal neurons. Hippocampus 22, 16591680 (2012).

[11] Xu, W. et al. Distinct neuronal coding schemes in memory revealed by selective erasure of fast synchronous synaptic transmission. Neuron 73, 990–1001 (2012).

[12] Payeur, A., Guerguiev, J., Zenke, F., Richards, B. A. & Naud, R. Burst-dependent synaptic plasticity can coordinate learning in hierarchical circuits. Nat. Neurosci. 24, 1010–1019 (2021).

[13] Wang, M. et al. Single-neuron representation of learned complex sounds in the auditory cortex. Nat. Commun. 11, 4361 (2020).

[14] Fujita, K., Kashimori, Y. & Kambara, T. Spatiotemporal burst coding for extracting features of spatiotemporally varying stimuli. Biol. Cybern. 97, 293–305 (2007).

[15] Miller, B. R., Walker, A. G., Barton, S. J. & Rebec, G. V. Dysregulated neuronal activity patterns implicate corticostriatal circuit dysfunction in multiple rodent models of Huntingtons disease. Front. Syst. Neurosci. 5, 26 (2011).

[16] Yang, Y. et al. Ketamine blocks bursting in the lateral habenula to rapidly relieve depression. Nature 554, 317–322 (2018).

[17] Lévy, P. Theorie de L’Addition des Variables Aleatoires. (Gauthier-Villars, Paris, 1954).

[18] Mandelbrot, B. The Fractal Geometry of Nature. (Freeman, New York, 1977).

[19] Abe, M. S. Functional advantages of Lévy walks emerging near a critical point. Proc. Natl. Acad. Sci. USA 117, 24336–24344 (2020).

[20] Viswanathan, G. M. et al. Optimizing the success of random searches. Nature 401, 911914 (1999).

[21] Bartumeus, F., Da Luz, M. G. E., Viswanathan, G. M. & Catalan, J. Animal search strategies: A quantitative random-walk analysis. Ecology 86, 30783087 (2005).

[22] Ott, A., Bouchaud, J., Langevin, D. & Urbach, W. Anomalous diffusion in living polymers: a genuine Levy flight? Phys. Rev. Lett. 65, 22012204 (1990).

[23] Brockmann, D., Hufnagel, L. & Geisel, T. The scaling laws of human travel. Nature 439, 462465 (2006).

[24] Huda, S. et al. Lévy-like movement patterns of metastatic cancer cells revealed in microfabricated systems and implicated in vivo. Nat. Commun. 9, 4539 (2018).

[25] Corral, A. Universal earthquake-occurrence jumps, correlations with time, and anomalous diffusion. Phys. Rev. Lett. 97, 178501 (2006).

[26] Barthelemy, P., Bertolotti, J. & Wiersma, D. A Lévy flight for light. Nature 453, 495498 (2008).

[27] Boccignone, G. & Ferraro, M. Modelling gaze shift as a constrained random walk. Physica A 331, 207–218 (2004).

[28] Sparks, D. L. & Barton E. J. Neural control of saccadic eye movements. Curr. Opin. Neurobiol. 3, 966–972 (1993).

[29] Kojima, Y. A neuronal process for adaptive control of primate saccadic system. Prog. Brain Res. 249, 169–181 (2019).

[30] Quinet, J., Schultz, K., May, P. J. & Gamlin, P. D. Neural control of rapid binocular eye movements: Saccade-vergence burst neurons. Proc. Natl. Acad. Sci. USA 117, 29123–29132 (2020).

[31] McNamee, D. C., Stachenfeld, K. L., Botvinick, M. M. & Gershman, S. J. Flexible modulation of sequence generation in the entorhinal-hippocampal system. Nat. Neurosci. 24, 851–862 (2021).

[32] Simonnet, J. & Brecht, M. Burst Firing and Spatial Coding in Subicular Principal Cells. J. Neurosci. 39, 3651–3662 (2019).

[33] Rhodes, T. & Turvey, M. T. Human memory retrieval as lévy foraging. Physica A 385, 255260 (2007).

[34] Costa, T., Boccignone, G., Cauda, F. & Ferraro, M. The Foraging Brain: Evidence of Lévy Dynamics in Brain Networks. PLoS One 11, e0161702 (2016).

[35] Patten, K. J., Greer, K., Likens, A. D., Amazeen, E. L. & Amazeen, P. G. The trajectory of thought: Heavy-tailed distributions in memory foraging promote efficiency. Mem. Cogn. 48, 772787 (2020).

[36] Sussillo, D. & Abbott, L. F. Generating coherent patterns of activity from chaotic neural networks. Neuron 63, 544–557 (2009).

[37] Sussillo, D., Churchland, M. M., Kaufman, M. T. & Shenoy, K. V. A neural network that finds a naturalistic solution for the production of muscle activity. Nat. Neurosci. 18, 1025–1033 (2015).

[38] Rajan, K., Harvey, C.D. & Tank, D. W. Recurrent Network Models of Sequence Generation and Memory. Neuron 90, 128–142 (2016).

[39] Enel, P., Procyk, E., Quilodran, R. & Dominey, P. F. Reservoir Computing Properties of Neural Dynamics in Prefrontal Cortex. PLoS Comput. Biol. 12, e1004967 (2016).

[40] Martín-Vázquez, G., Asabuki, T., Isomura, Y. & Fukai, T. Learning Task-Related Activities From Independent Local-Field-Potential Components Across Motor Cortex Layers. Front. Neurosci. 12, 429 (2018).

[41] Denève, S. & Machens, C. K. Efficient codes and balanced networks. Nat. Neurosci. 19, 375–382 (2016).

[42] Abbott, L., DePasquale, B. & Memmesheimer, R. M. Building functional networks of spiking model neurons. Nat. Neurosci. 19, 350355 (2016).

[43] Bellec, G. et al. A solution to the learning dilemma for recurrent networks of spiking neurons. Nat. Commun. 11, 3625 (2020).

[44] Thalmeier, D., Uhlmann, M., Kappen, H. J. & Memmesheimer, R-M. Learning Universal Computations with Spikes. PLoS Comput. Biol. 12, e1004895 (2016).

[45] Kim, C. M. & Chow, C. C. Learning recurrent dynamics in spiking networks. eLife 7, e37124 (2018).

[46] Nicola, W. & Clopath, C. Supervised learning in spiking neural networks with FORCE training. Nat. Commun. 8, 2208 (2017).

[47] Izhikevich, E. M. Simple model of spiking neurons. IEEE Trans. Neural Netw. 14,1569–1572 (2003).

[48] Optican, L. M. & Pretegiani, E. What stops a saccade? Phil. Trans. R. Soc. B 372, 20160194 (2017).

[49] Epsztein, J., Brecht, M. & Lee, A. K. Intracellular determinants of hippocampal CA1 place and silent cell activity in a novel environment. Neuron 70, 109120 (2011).

